# Ribosomal protein L24 modulates mammalian microRNA processing and transfer RNA fragment production

**DOI:** 10.1101/2023.05.03.539194

**Authors:** Yonat Tzur, Serafima Dubnov, Nimrod Madrer, Adi Bar, Bettina Nadorp, Nibha Mishra, Paul Heppenstall, Estelle R Bennett, David S Greenberg, Katarzyna Winek, Hermona Soreq

**Author notes:** Co-corresponding authors: Katarzyna Winek and Hermona Soreq; Katarzyna Winek, Hermona Soreq, Edmond J. Safra Campus Givat Ram, 91904 Jerusalem, Israel. CSL Seqirus, 225 Wyman Street, Waltham, MA 02451.

## Abstract

The evolutionary mechanism(s) underlying the expression of novel microRNAs (miRs) are still elusive. To explore this issue, we studied the expression of intronic primate-specific hsa-miR-608, located in the Semaphorin 4G (SEMA4G) gene. Engineered ‘humanized’ mice carrying human miR-608 flanked by 250 bp in the murine Sema4g gene expressed miR-608 in several tissues. Moreover, miR-608 flanked by shortened fragments of its human genome region elevated miR-608 levels by 100-fold in murine and human-originated cells, identifying the 150 nucleotides 5’ to pre-miR-608 as an active promoter. Surprisingly, pulldown of this 5’ sequence revealed tight interaction with ribosomal protein L24 (RPL24), which inhibited miR-608 expression. Furthermore, RPL24 depletion altered the levels of 22 miRs, and we discovered that direct interaction of RPL24 with DDX5, a component of the large microprocessor complex, inhibits pri-miR processing. Moreover, RPL24 depletion resulted in Angiogenin (ANG)-mediated production of 5’-half tRFs in human cells, and altered plant tRF profiles. Expanding previous reports that RPL24 regulates miR processing in Arabidopsis thaliana, we implicate RPL24 in an evolutionarily-conserved regulation of miR processing and tRF production.

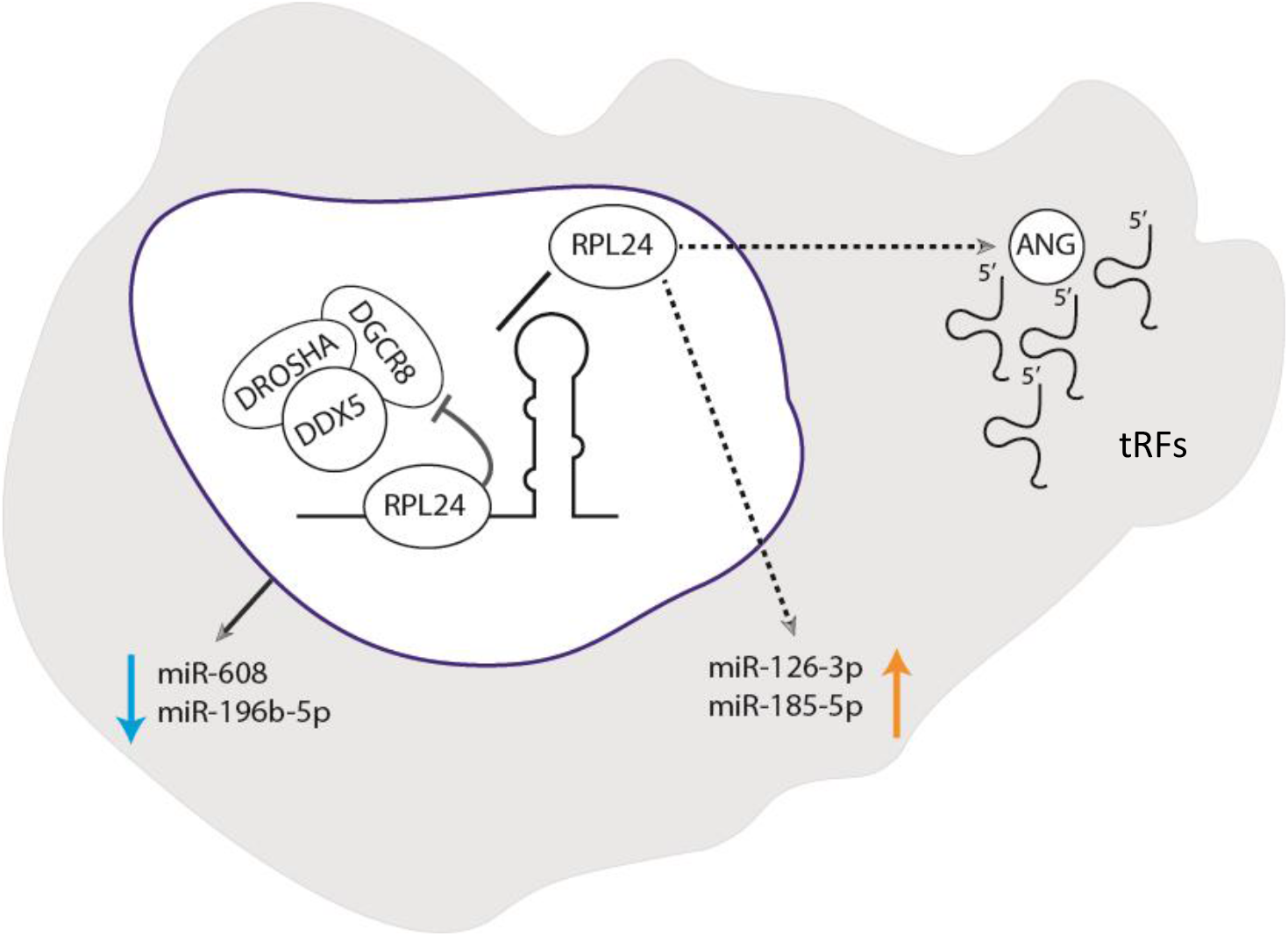

## Introduction

Mammalian gene expression is modulated by microRNAs (miRs), small (∼22 nucleotides) non-coding RNAs which act by binding to mRNA transcripts that contain complementary sequence motifs and silencing them post-transcriptionally [1]. Importantly, a large fraction of the known human miRs have been evolutionarily incorporated into the primate genome [2], but neither the mechanism(s) enabling their expression nor their global biological impact are known. In metazoans, RNA polymerase II transcribes the majority of primary miRs (pri-miRs), followed by nuclear processing in the microprocessor complex. This complex is comprised of the core components nuclear RNase III DROSHA and the double-stranded RNA-binding protein DGCR8, and includes additional auxiliary factors such as the RNA helicases DDX5 (p68) and DDX17 (p72) [3]. The resulting pre-miRs are transported to the cytoplasm by exportin 5 [4], where they are further processed by the RNase III Dicer. Mature miRs are then loaded onto a multi-protein RNA-induced silencing complex (RISC) that inhibits translation and/or promotes degradation of the target mRNA transcripts [5]. However, much remains unclear about the mechanism of microprocessor recognition and cleavage of the pri-miRs. Further, the recently re-discovered transfer RNA fragments (tRFs), 16-50 nucleotides long, have diverse biological functions [6], among them an active contribution to the cholinergic anti-inflammatory impact [7] and regulation of gene expression [8]. Nonetheless, the mechanisms controlling tRF production are incompletely understood.

The ‘basal’ junction between the single-stranded RNA and the double-stranded stem RNA of pre-miRs, and the size of the terminal loop of pri-miRs are all crucial for microprocessor recognition and processing [9, 10]. In addition, several sequence elements such as the UG and CNNC motifs residing in the basal region of pri-miRs and the UGUG motif in the terminal loop are involved in pri-miR processing [11, 12]. However, only some pri-miRs carry these motifs, suggesting that other regulatory elements are involved as well. In this context, hsa-miR-608, located in the third intron of the primate SEMA4G gene, has been hypothesized to share a common promoter with its host gene [13, 14], However, the mechanisms underlying its expression remained elusive, as pri-miR-608 contains none of the motifs described above.

At the functional level, in humans miR-608 targets the acetylcholine hydrolyzing enzyme acetylcholinesterase (AChE), the inflammatory cytokine interleukin-6 (IL-6) and the cell division cycle 42 (CDC42) Rho GTPase, indicating relevance for the cholinergic blockade of inflammation [15] and reaction to stressful stimuli [16]. However, AChE’s recognition by miR-608 is interrupted by the clinically significant [12] rs17228616 single-nucleotide polymorphism (SNP) found in the 3’-untranslated region (3’-UTR) of the AChE gene. Carriers of the minor, less abundant allele of this SNP show reduced miR-608 blockade of AChE which results in lower cholinergic tone [16]. This is accompanied by elevated blood pressure, increased inflammatory biomarkers, and prefrontal cortex blockade of amygdala stress reactions [16, 17]. In comparison, the rs4919510 SNP resides in the miR-608 gene itself and carriers of its minor allele show reduced miR-608 levels, limited miR-608 regulation of AChE and other targets, and protection from sepsis following head injury [18]. Given the importance of miR-608 contribution to the regulation of the cholinergic tone, we sought regulatory elements facilitating miR-608 expression and contributing to its impact on AChE expression and human health and wellbeing.

## Results

### Cis-sequences regulate miR-608 expression

To explore and characterize the molecular regulation of miR-608 we established a ‘humanized’ hsa-miR-608 mouse by inserting the pre-miR-608, flanked by 250 bases of intronic sequences from its human host gene SEMA4G, into the third intron of the mouse Sema4g gene (Figure 1A). Quantifying miR-608 levels in various brain areas as well as peripheral tissues from miR-608 knock-in (KI) mice revealed miR-608 expression in the hypothalamus and cerebellum (Figure 1B) as well as in the intestine, liver, and kidney, with peripheral tissues showing higher levels than brain tissues, and no difference between female and male mice.. Nevertheless, the transgenic mice maintained unchanged motor and anxiety traits as measured by rotarod, elevated plus maze and open field tests (Supplementary Figure 1). Interestingly, miR-608 levels were below detection in the hippocampus, medial-prefrontal cortex, and heart, indicating tissue-specific regulation of miR-608 expression in these mice.

**Figure 1:**
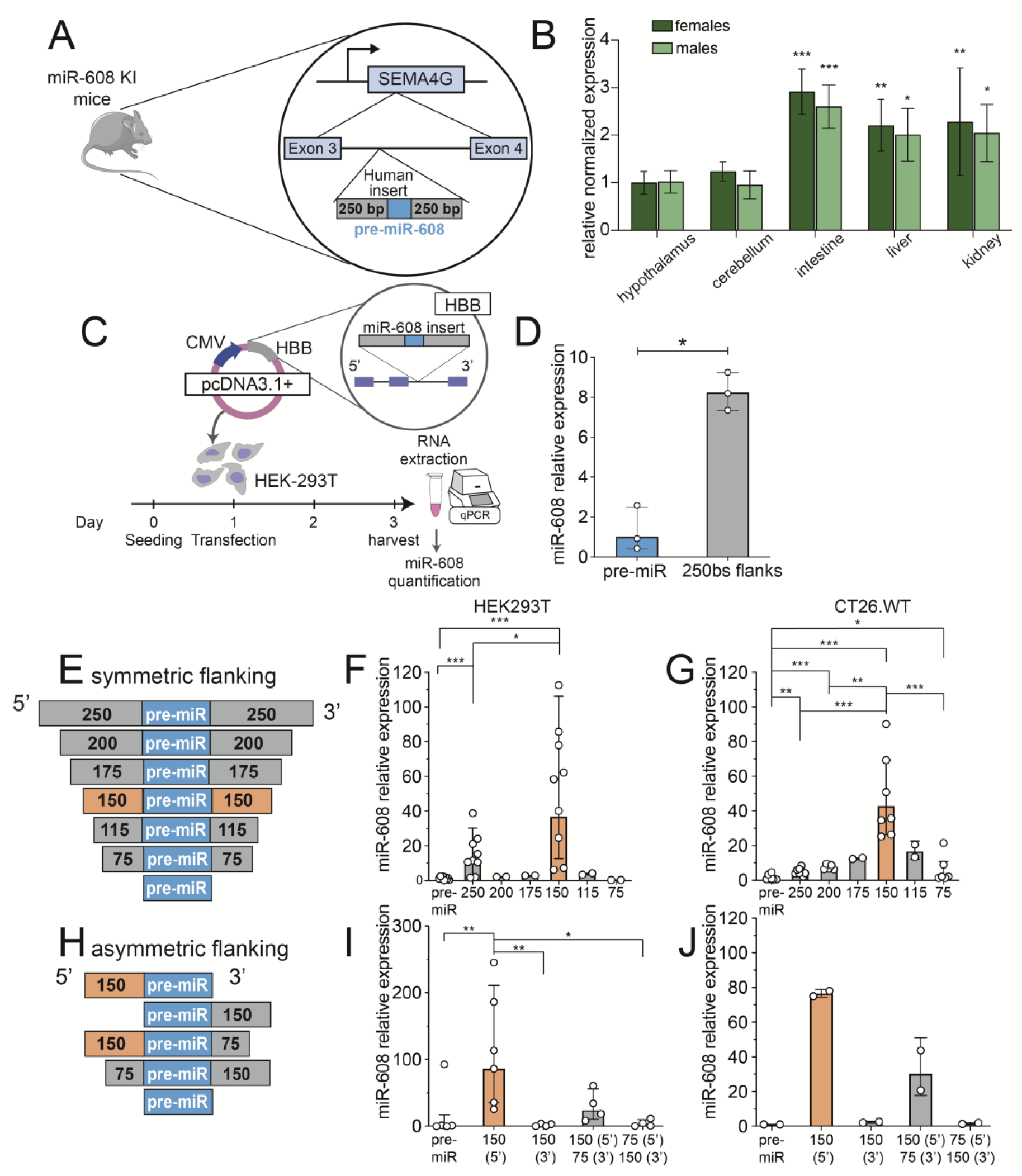
miR-608 cis-sequences regulate its expression in engineered KI mice and cultured cells. **A**. Structure of the mouse construct: miR-608 transgenic ‘humanized’ mice were established by inserting hsa-pre-mir-608 into the third intron of the mouse Sema4g gene, with 250 flanking bases at each side. **B**. miR-608 levels in brain and peripheral tissues of female and male miR-608 KI mice, determined by RT-qPCR. miR-608 levels were normalized to those of female hypothalamus (showing the lowest expression); two-way ANOVA with Dunnet’s multiple comparisons correction, ±SD, n=5 per group. **C**. Experimental design: HEK293T and CT26.WT cells were seeded and 24 hours later transfected with pcDNA3.1+ plasmid containing miR-608 inserted into the second intron of HBB and flanked by native sequences of varying lengths. 48 hours later cells were harvested, RNA extracted, and miR-608 levels quantified. **D**. Symmetric bidirectional 250 base extension of the miR-608 stem-loop altered its levels in HEK293T cells; bar graph ±SD, p=0.0163, unpaired t-test. **E**. Constructs of pre-miR-608 flanked by symmetric sequences ranging from 75 to 250 bases. **F**. Levels of miR-608 expressed from these constructs in HEK293T cells. The 150 bases both upstream and downstream are critical for miR-608 expression. **G**. The 150-base symmetrical flanks are also critical for expression in CT26.WT cells. **H**. Constructs of pre-miR-608 flanked by asymmetric sequences ranging from 75 to 150 bases. **I**. Induced levels of miR-608 expressed from these constructs in HEK293T cells were highest under control of the upstream 150 base sequence. **J**. A similar effect is seen in CT26.WT cells transfected with these constructs. All experiments were performed in duplicate or triplicate and miR-608 levels were measured using Taqman RT-qPCR with RNU6B and snoRNA135 as normalizing genes. Results are shown relative to levels of pre-miR-608 with no flanking sequences. In all panels * p <0.05, ** p <0.01, *** p <0.001. In panels F, G, I, bar-graph ± SD, one-way ANOVA with Tukey’s multiple comparisons test.

Given that miR-608 is primate-specific, we speculated that its expression in a non-primate organism was facilitated by the flanking intronic sequences from its human host gene. To identify flanking sequences critical for miR-608 expression we created a series of constructs containing pre-miR-608 surrounded by different lengths of the native flanking bases. We then inserted these constructs into the second intron of the human hemoglobin subunit beta (HBB) gene that we cloned into the pcDNA3.1+ plasmid. We transfected both human (HEK293T) and murine (CT26.WT) cell lines with the different constructs and quantified mature miR-608 levels 48 hours after transfection (Figure 1C). Since basal miR-608 levels in HEK293T cells were below detection threshold (Sup. Fig 2A) (and no basal expression is expected in the non-primate CT26.WT cells), any detected miR-608 could be attributed solely to the transfected constructs.

**Figure 2:**
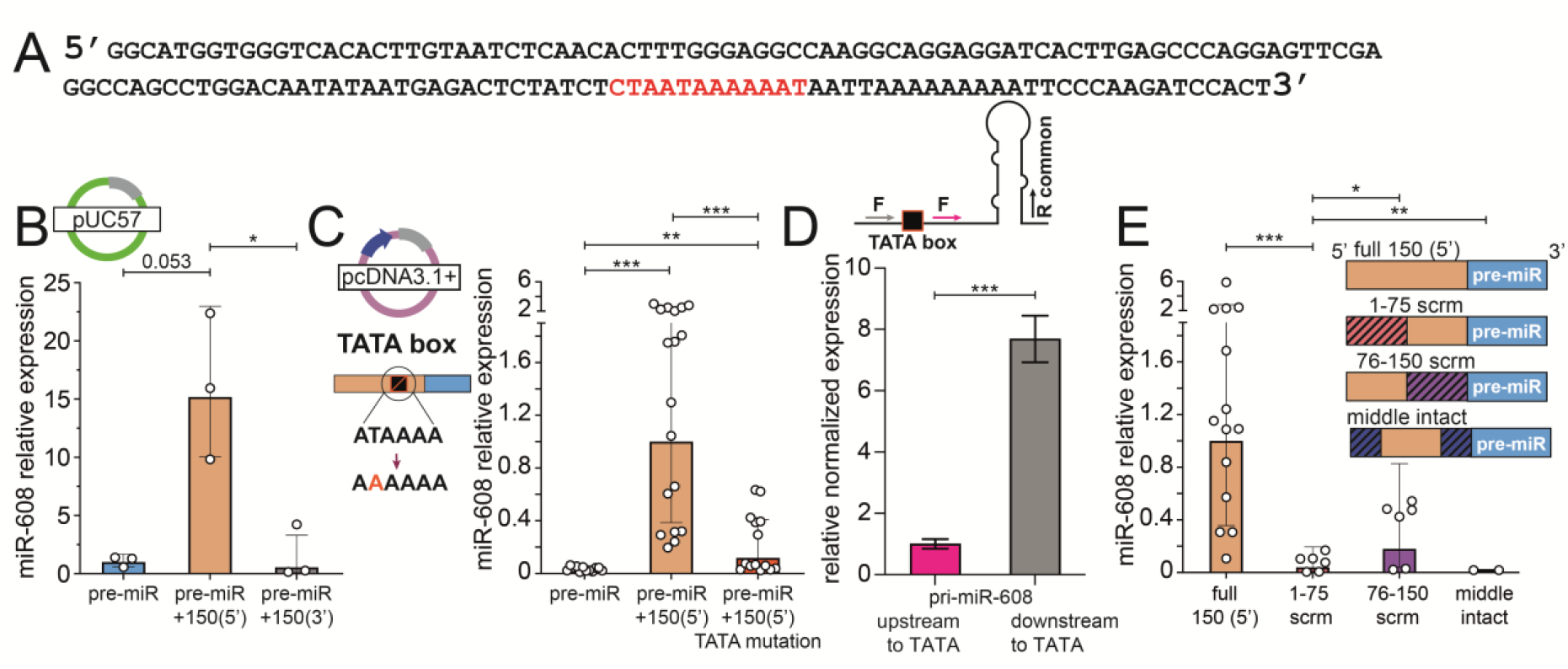
The 5’ 150 bases contain a TATA box enabling miR-608 expression. **A**. The 150 bases 5’ upstream to pre-miR-608 contain a TATA box at position 111 (red). **B**. miR-608 levels in HEK293T cells transfected with pUC57 quantified by Taqman RT-qPCR, showing that the TATA box drives expression from the promoter-less bacterial vector pUC57. **C**. miR-608 levels in HEK293T cells transfected with the mammalian vector pcDNA3.1+ containing the TATA box T → A point mutation are reduced as compared to non-mutated TATA box. **D**. Higher levels of pri-miR-608 transcripts (expressed from pcDNA3.1+) and measured by RT-qPCR using forward primers positioned downstream (F2) vs upstream (F1) to the predicted TATA box; the reverse primer (R) is common. **E**. Scrambling parts of the 5’ 150 bases upstream to the pre-miR-608 sequence decreased the levels of mature miR-608. Hatching represents the scrambled area in the sequence. Taqman RT-qPCR quantification of miR-608 yielded results relative to the intact 150 base sequence and normalized to RNU6B. In all panels * p <0.05, ** p <0.01, *** p <0.001. In panel D unpaired t-test, in all other panels one-way ANOVA with Tukey’s correction for multiple comparisons, bar-plot ± SD.

Transfecting cells with pre-miR-608 flanked by 250 bases at both the 5’ and 3’ ends resulted in miR-608 levels 8-fold higher than in cells transfected with the pre-miR-608 lacking flanking sequences, indicating that the flanking sequences contain sufficient elements to enable miR-608 expression (Figure 1D). To challenge this concept, we introduced symmetric flanks of gradually shortened lengths at both the 5’ and 3’ ends of pre-miR-608 and compared their expression efficacy to that of the pre-miR-608 construct with no flanking regions. Constructs with flanking sequences of 150 bases at both the 5’ and 3’ sides increased miR-608 expression by over ∼40-fold in both HEK293T and CT26.WT cells (Figure 1E-G), indicating the possible presence of binding motifs for transcription activators within these sequences. In contrast, extending the symmetric flanking beyond those 150 bases greatly decreased pre-miR-608 levels, possibly reflecting the presence of binding motifs for transcription suppressors in the sequences 150-250 bases upstream and/or downstream of pre-miR-608. To identify sequences indispensable for miR-608 expression we expressed pre-miR-608 constructs with asymmetric or lacking flanks (1H). Most asymmetric combinations of flanking sequences showed low expression in both cell lines with the exception of the 5’ 150-base flanking sequence alone that was sufficient to potentiate miR-608 expression ∼100-fold in HEK293T and ∼80-fold in CT26.WT cells, such that the absence of the 3’ sequence caused an increase of roughly 2-fold in miR-608 expression in both cell lines (Figure 1H-J).

### The 5’ sequence includes a TATA box enabling miR-608 expression

As the 5’ 150-base sequence showed the most pronounced effect on miR-608 expression, we employed the “YAPP Eukaryotic Core Promoter Predictor” bioinformatic tool (http://www.bioinformatics.org/yapp/cgi-bin/yapp.cgi) to search for predicted promoter features and activator binding sites in this region. We identified a TATA box (CTAATAAAAAAT) at position 111 in the sequence (Figure 2A) with high certainty (score of 0.89). This TATA box could potentially serve as a core promoter sufficient to drive miR-608 expression in a promoter-less vector. To test this prediction, we inserted pre-miR-608 and its 5’ 150 bases into the pUC57 bacterial vector which is devoid of a mammalian promoter. Supporting our prediction, when transfected into HEK293T and CT26.WT cells, the 5’ 150 bases elevated miR-608 levels about 15-fold over the pre-miR alone or the construct including the 3’ 150 bases. Thus, even without an external promoter both cell lines expressed miR-608 (Figure 2B, Sup. Fig 2B). To further assess the importance of the TATA box we mutated thymine at position 5 (marked in bold-face in Figure 2A) to adenine and transfected HEK293T cells with this construct in pcDNA3.1+, not pUC57, to better measure changes in expression from a base-line of 100-fold vs. 15-fold over pre-miR-608 (Figure 1I vs. Fig 2B). Levels of miR-608 were reduced by over 80%, confirming the indispensability of the TATA box in driving miR-608 expression (Figure 2C).

The impact of the TATA core promoter on miR-608 expression was further tested by comparing pri-miR-608 levels that reflect transcription initiated from the CMV promoter (using a forward primer upstream to the TATA box) vs. pri-miR-608 levels that reflect transcription initiated from the TATA box itself (using a forward primer downstream to the TATA box) (Figure 2D, upper section). Predictably, the level of pri-miR-608 transcripts initiated by the TATA core promoter were close to 8-fold higher than the levels initiated by the CMV promoter (Figure 2D, lower section). In other words, roughly 85% of miR-608 is transcribed from the core promoter. This agrees with the results shown in Figure 2C, whereby TATA box mutation reduced miR-608 by over 80%. Interestingly, partially scrambling stretches of 75 nucleotides in these 150 bases abolished (1-75) or greatly reduced (76-150) miR-608 expression, suggesting that expression is affected by additional factors such as secondary structure or the location of important regulatory binding sites in this sequence (Figure 2E, left and right panels).

### RPL24 affects mammalian miR biogenesis

Numerous proteins impact the processing and expression levels of miRs [19, 20]. To identify proteins that interact with the 5’ 150 bases that control miR-608 levels, we performed a pull-down assay in which we incubated a lysate of HEK293T cells with a biotinylated oligonucleotide comprised of the 5’ 150-base sequence. The proteins bound to the sequence were then isolated using streptavidin-coated magnetic beads (sup. Fig 3A) and identified by mass spectrometry (MS). This revealed the ribosomal protein RPL24 as the most enriched protein bound to the 5’ 150-base sequence (Figure 3A). RPL24, a member of the L24E family of ribosomal proteins, is a component of the 60S ribosomal subunit and contributes to ribosome assembly and translational processes in the cytoplasm [21]. In *Arabidopsis thaliana*, a portion of STV1 — the plant homolog of RPL24 — localizes to the nucleus and affects the level of various miRs by binding a short 5’ sequence on the pri-miR and influencing their interaction with HYL1, a component of the plant microprocessor [22].

**Figure 3:**
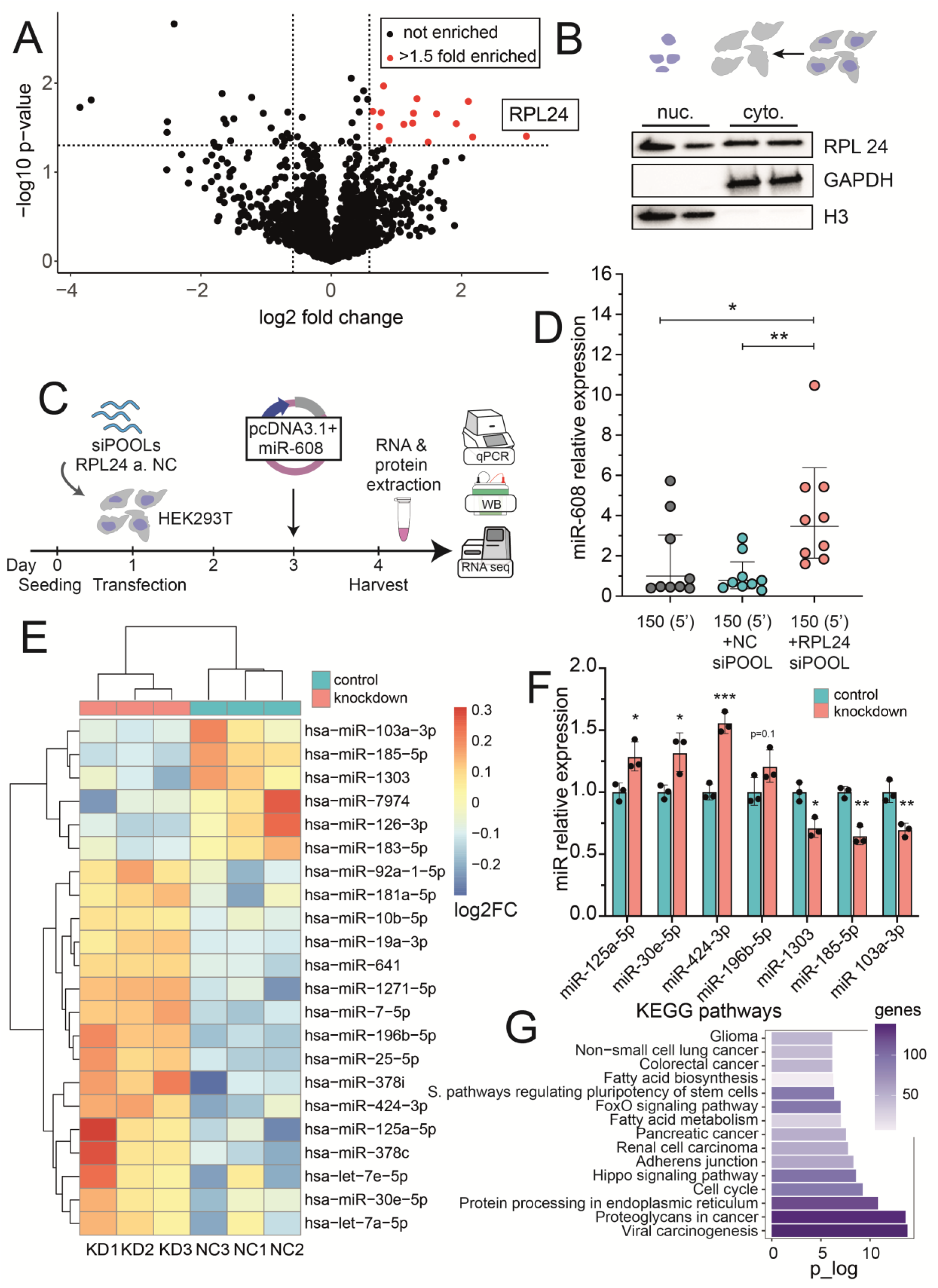
RPL24 KD alters the levels of diverse miRs in addition to miR-608. **A**. Pulldown analysis: mass spectrometry identified 15 proteins bound to the 5’ 150 base-upstream sequence with p<0.05 and enrichment >1.5-fold, the most enriched being RPL24 (p = 0.039, fold enrichment = 8). **B**. Immunoblot of subcellular fractions identified RPL24 in both the nuclear and the cytoplasmic fractions. Fraction purity was validated with GAPDH, a cytoplasmic marker, and H3, a nuclear marker; each was localized solely to its expected compartment. **C**. Experimental design: HEK293T cells were seeded, transfected with 50nM of siPOOLs targeting RPL24 or a non-targeting negative control pool (NC) 24 hours later, then transfected with pcDNA3.1+ containing miR-608 48 hours later. RNA and proteins were extracted 24 hours post-transfection. **D**. RPL24 depletion elevates miR-608 levels as shown by Taqman RT-qPCR quantification of miR-608 levels, with results normalized to RNU6B and relative to NC siPOOLs; one-way ANOVA with Tukey’s correction for multiple comparisons ±SD. **E**. Heatmap showing the 22 DE miRs elevated or decreased after RPL24 knockdown, analyzed by DESeq2, adjusted p-value <0.05, Benjamini-Hochberg correction [27]. See Sup. table 1 for full information on the DE miRs. **F**. RT-qPCR validation confirming the RNA-seq results with data relative to the NC and normalized to SNORD47 and SNORD48 l; unpaired t-test for each miR± SD. **G**. KEGG analysis showing the 15 most enriched pathways for targets of miRs regulated by RPL24. In all panels * p <0.05, ** p <0.01, *** p <0.001.

To investigate whether RPL24 may also contribute to miR processing in mammalian cells, we first sought to determine if RPL24 is present in the nuclear compartment in which pri-miRs are transcribed and processed [5]. Using a standard fractionation protocol in HEK293T cells (see Materials and Methods) we confirmed that RPL24 is indeed found in both this nuclear fraction and the expected cytoplasmic fraction (Figure 3B). To determine if this location indicates a possible role for RPL24 in regulating miR-608 expression in human cells we administered siRNAs targeting RPL24 (siPOOLs) to knock-down RPL24 in HEK293T cells (Figure 3C). RPL24 RNA levels were decreased by ∼80% and protein levels by ∼60% (Sup. Fig 3B-D). In contrast, the levels of mature miR-608 (Figure 3D) were increased over 3-fold as compared to negative control (NC) siPOOLs, thus suggesting that RPL24 binding to the 5’ sequence on pri-miR-608 inhibits the expression of the mature miR-608 in mammalian cells.

To examine the global impact of RPL24 on miR processing we performed a separate RPL24 KD experiment in HEK293T cells followed by small RNA sequencing. We were able to identify 22 differentially expressed (DE) miRs, 16 of which were increased and 6 decreased when compared to NC (Figure 3E, Sup. table 1). As HEK293T cells do not express miR-608, it was not detected in the sequencing dataset. RT-qPCR-quantification validated the sequencing data and confirmed significant changes in the levels of 6 out of 11 tested DE miRs; miR-125a-5p, miR-30e-5p and miR-424-3p were upregulated whereas miR-1303, miR-185-5p and miR-103a-3p were downregulated after RPL24 KD (Figure 3F). These data support the general involvement of RPL24 in miR biogenesis in mammalian cells. The targets of the above DE miRs were identified (using DIANA Tools, [23]) and KEGG pathway analysis detected a marked enrichment in cancer-related pathways Figure 3G). Thus, while the role of RPL24 in tumorigenesis has so far been attributed to its involvement in translation [24-26], our current findings add RPL24 regulation of a specific set of miRs as potentially contributing to various cancers.

### RPL24 KD induces production of 5’-half tRFs

The influence of RPL24 and its plant homolog STV1 on miR biogenesis raised the question whether additional small non-coding RNA (sncRNA) classes, such as transfer RNA fragments (tRFs), are also affected in an evolutionarily conserved manner between mammalian cells and *Arabidopsis thaliana* (Figure 4A). Using the MINTmap tool [28], we searched for DE tRFs by analyzing the RNA-seq dataset obtained following RPL24 KD in HEK293T cells. Principal component analysis (PCA) completely separated the RPL24 KD and the NC groups (Figure 4B) and we found altered levels of 20 nuclear genome-originated, 31-35 base long tRFs, 19 of which were increased following RPL24 KD (Figure 4C). Of these, 17 were 5’-half tRFs, with seven of them originating from histidine and eight from glutamine tRNAs (Figure 4D, Sup. table 2). Moreover, electrophoretic size selection followed by RT-qPCR quantification confirmed the specific upregulation of the DE tRFs originated from full-length histidine tRNAs (materials and methods; Figure 4E). 5’-half tRFs are produced in the cytoplasm by angiogenin (ANG) in response to cellular stress and facilitate cell survival by promoting the formation of stress granules and inhibiting apoptosis [6, 29, 30]. To test if ANG is involved in generation of the specific tRFs upregulated in our RPL24 KD experiment, we analyzed a dataset by Su et al. (GEO accession number GSE130764, [29]). Supporting our prediction, ANG overexpression in HEK293T cells induced expression of a wide range of tRFs, with the most significant of them identical to those upregulated following RPL24 KD (Figure 4F). Therefore, that RPL24 KD results in ANG-mediated production of 5’-half tRFs may reflect the cellular stress and translation inhibition caused by RPL24 depletion.

**Figure 4:**
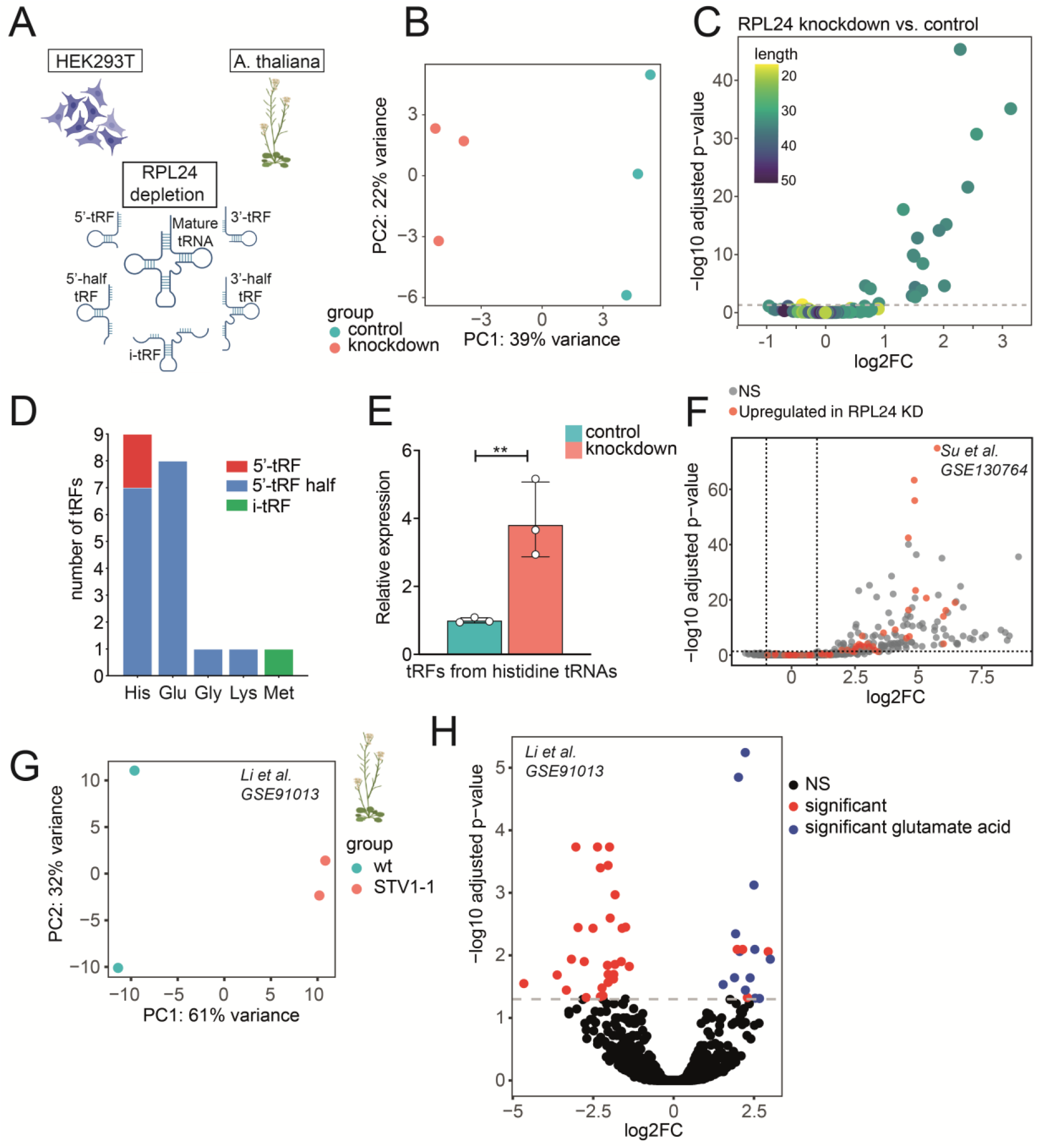
RPL24 KD results in ANG-mediated production of 5’-half tRFs. **A**. The effect of RPL24 depletion on tRF level and type was tested in HEK293T cells and analyzed in *Arabidopsis thaliana*. **B**. PCA of DE tRFs in HEK293T cells after RPL24 KD shows that samples are separated by treatment. **C**. Volcano plot of DE tRFs following RPL24 KD revealed 19 increased and 1 decreased tRFs (p-adj < 0.05). **D**. Bar graph showing the type and amino acid tRNA origin of the DE tRFs. **E**. RT-qPCR validation of the tRFs derived from his-tRNAs, following size selection of small RNA. Results are normalized to miR-145-5p and miR-15a-5p, p =0.0014, unpaired t-test, ± SD, ** p <0.01. **F**. Volcano plot of DE tRFs following ANG overexpression in HEK293T cells. Red and grey symbols denote tRFs significantly upregulated following RPL24 KD and non-significantly changed following RPL24 KD, respectively (p-adj < 0.05). **G**. PCA of DE tRFs in *Arabidopsis thaliana* showing separation of WT and STV1-1 samples. **H**. Volcano plot of DE tRFs in *Arabidopsis*; Red, blue and black symbols denote tRFs originating from all amino acids, originating from glutamic acid or non-significantly altered tRFs.

To further address the impact of the evolutionary conservation of RPL24/STV1 on tRF levels we analyzed a small RNA-seq from *Arabidopsis thaliana* (GEO accession number GSE91013, [22]), composed of 2 wild type (WT) and 2 mutant (STV1-1) samples, using tRFanalyzer [31]. Similar to our finding in HEK293Tcells, PCA completely separated the WT from the STV1-1 group (Figure 4G). 45 tRFs were significantly DE between the groups, with 28 decreased and 17 increased (Figure 4H, sup. table 3). Importantly, most of the increased tRFs originated from glutamic acid-carrying tRNAs, similar to our findings in mammalian HEK293T following RPL24 KD. However, unlike RPL24 KD in mammalian cells, in the mutant plants the majority of tRFs are decreased with no specific elevation of 5’-halves. This may reflect adaptation to the chronic stress resulting from lack of STV1 vs. the acute stress caused by RPL24 depletion in HEK293T cells.

### RPL24 interacts with DDX5 to inhibit pri-miR processing

To further investigate the nature of RPL24 involvement in miR biogenesis and tRF production we immunoprecipitated (IP) a FLAG-tagged RPL24 followed by MS analysis to identify interacting proteins. To avoid masking of RPL24’s nuclear interactors by cytoplasmic partners we performed separate IPs for the nuclear and cytoplasmic fractions (Figure 5A). Immuno-blotting confirmed the marked enrichment of FLAG-RPL24 in the pellet (Figure 5B) and interacting partners were identified in subsequent MS analysis. Over 100 proteins were found in each of the compartments (Figure 5C-D, sup table 4 and 5), with over half of them shared (Figure 5E). In agreement with our hypothesis that RPL24 indirectly upregulates the levels of ANG-produced tRF halves, no interaction of FLAG-RPL24 with ANG was detected in the cytoplasmic fraction. Importantly, the RNA helicase DDX5 was enriched in the nuclear fraction (4-fold, p=0.003 Figure 5C), demonstrating potential RPL24 interaction with the microprocessor. Immunoblot analysis of FLAG-RPL24 IP in an independent experiment confirmed this interaction (Figure 5F).

**Figure 5:**
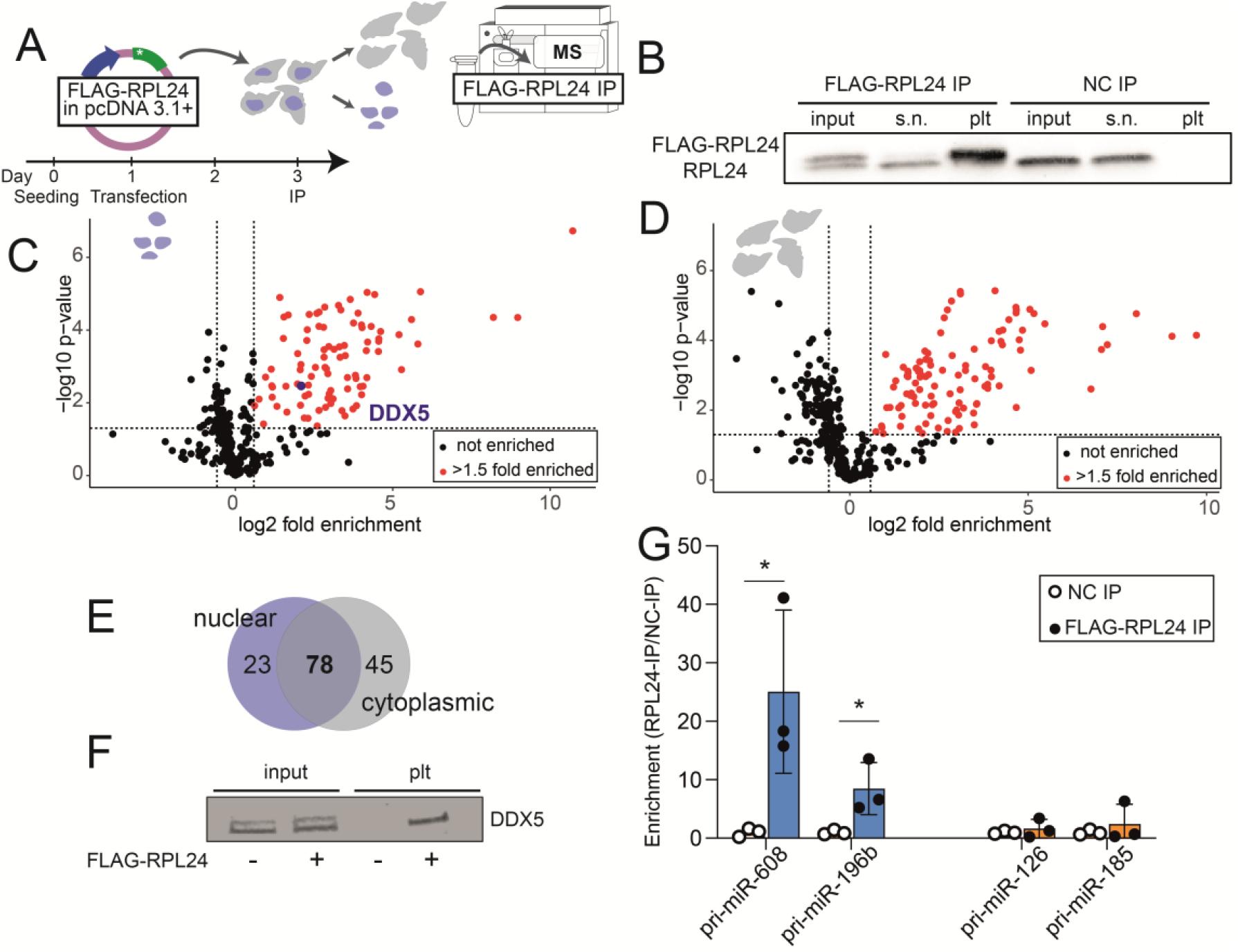
RPL24 inhibits pri-miR processing through direct interaction with DDX5 and pri-miR-608. **A**. Experimental design: HEK293T cells were seeded and 24 hours later transfected with pcDNA3.1+ vector containing FLAG-RPL24 or no insert as control. Cells were harvested 48 hours after transfection, fractionated, and nuclear and cytoplasmic fractions immunoprecipitated separately with an antibody against the FLAG-tag, with subsequent MS analysis. **B**. Immunoblot of nuclear fraction from cells transfected with a pcDNA3.1+ vector containing FLAG-RPL24 insert or empty pcDNA3.1+ vector as control. 2.5% of input lysate was loaded per lane, 2.5% of supernatant, and 20% of pellet. A double band is observed in the input lane; the upper band is the FLAG-RPL24 (∼24kDa) and the lower band is the endogenous RPL24 (∼23kDa). **C, D**. Volcano plot presenting the proteins enriched in FLAG-RPL24 IP compared to control, in the nuclear and cytoplasmic fractions. Black and red symbols denote non-significant proteins and proteins with p<0.05 and enrichment >1.5-fold. **E**. Venn diagram showing proteins bound to RPL24 in cytoplasmic and nuclear fractions, with common and specific proteins. **F**. Immunoblotted DDX5 from the nuclear fraction of HEK293T cells transfected with pcDNA3.1+ vector containing or lacking (control) a FLAG-RPL24 insert, confirming RPL24-DDX5 interaction. 2.5% of input lysate and 90% of pellet were loaded per lane. **G**. RT-qPCR quantification of pri-miRs in nuclear pellet samples of FLAG-RPL24 IP. Pri-miR-608 and pri-miR-196b are enriched, p =0.04 and p =0.045, respectively, but pri-miR-185 and pri-miR-126 are not expressed. Each pellet sample was normalized to its corresponding input sample and fold enrichment is determined as FLAG-RPL24 IP/NC IP.

To affirm that RPL24 executes its inhibiting effect on miR levels by binding to their pri-miRs, we immunoprecipitated RPL24, extracted RNA from the nuclear pellet fraction, and quantified the levels of pri-miR-608 and pri-miR-196b, the two most statistically significant miRs elevated by RPL24 KD. Both were enriched, indicating direct interaction between the protein and these pri-miRs (Figure 5G, sup table 1). In contrast, levels of pri-miR-185 and pri-miR-126, the two most statistically significant miRs reduced by RPL24 KD (sup table 1), did not reveal any enrichment in the IP pellet (Figure 5G), indicating a non-direct effect of RPL24 on these pri-miRs. Together, these findings suggest that direct interaction of RPL24 with pri-miRs and with the microprocessor via DDX5 inhibits pri-miR processing. However, miRs that were downregulated following RPL24 KD did not interact with RPL24 in the nucleus, and their levels are most likely modified by other factors.

## Discussion

To explore the mechanisms controlling expression of the primate-specific miR-608, we created a transgenic mouse model expressing miR-608 flanked by 250 nucleotide-long regions and integrated into the 3rd intron of the mouse Sema4g gene. We discovered hsa-miR-608 expression in several tissues of the engineered mice and revealed that the human intronic flanking sequences are functionally involved in regulating its expression. Specifically, we identified a 150 nucleotide-long sequence 5’ to pre-miR-608 which elevated miR-608 levels 100-fold in transfection tests of miR-608 flanked by serially shortened regions from its human genome location. Moreover, we identified a TATA box within this sequence that acts as a core promoter to induce miR-608 expression independently of its host gene promoter. This is compatible with reports that intronic miRs that are co-expressed from the host promoter show low evolutionary conservation [32]. Nevertheless, independent expression of newer miRs such as the primate-specific miR-608 may confer tighter regulation and availability to function regardless of their host gene expression.

Pulldown experiments identified the involvement of RPL24 in regulating mammalian miR levels. RPL24 is a key component of the ribosomal 60S subunit that is common to diverse organisms from archaea to eukaryotes. It operates as a translation factor which incorporates into the large ribosomal subunit and facilitates interaction between the large and small ribosomal subunits [21]. In mice homozygous Rpl24 deficiency is lethal and heterozygous mice are viable but develop various abnormalities [33]. In *Arabidopsis thaliana*, mutated plants lacking the RPL24 homolog short valve 1 (STV1) demonstrate altered levels of several miRs compared to WT plants. STV1 localizes to the nucleus where it binds a short 5’ sequence of pri-miRs and promotes their cleavage by facilitating their interaction with the HYPONASTIC LEAVES 1 (HYL1) protein, a component of the plant miRNA processing complex [22].

The ribosomal protein STV1 (RPL24) thus participates in the biogenesis of plant miRs, but our findings reveal that its role is evolutionarily conserved in miR processing in animals, including primates. We have shown for the first time that RPL24 is found not only in the cytoplasm but also in the nucleus of mammalian cells. In this cellular compartment it binds a 5’ 150-nucleotide-long sequence upstream to pre-miR-608 and inhibits miR-608 expression. Demonstrating a broader role of RPL24 in miR biogenesis, our small RNA-seq analysis following in vitro RPL24 depletion revealed altered levels of 22 miRs, suggesting that, also in mammals, RPL24 is actively involved in fine-tuning the levels of numerous miRs. Thus, our findings suggest an evolutionary conservation of RPL24’s role in miR biogenesis. Intriguingly, several eukaryotic ribosomal proteins (e. g. RPL4, RPS4, 6, 7, 9 and 14) are imported from the cytoplasm to the nucleolus where they participate in the assembly of ribosomal subunits that are then exported to the cytoplasm [34, 35]. Other ribosomal proteins have extra-ribosomal functions, including RPL11 which binds the MYC protein and inhibits the transcriptional activation of MYC-targeted genes [36] and RPS3 which can act as a caspase-dependent inducer of apoptosis and a DNA repair endonuclease [37, 38]. However, to our knowledge, none of these nuclear ribosomal proteins were shown to participate in pri-miR processing in mammalian cells.

Importantly, our findings also show that RPL24 depletion results in elevated levels of a recently re-discovered family of noncoding RNAs, the tRFs. Recent studies reveal that tRFs participate in regulating many biological processes, including gene expression, translation and RNA processing and stability [6, 39-41]. Some tRFs function like miRs, interacting with AGO proteins to silence their target mRNA [7, 42], while others were shown to regulate the levels of ribosomal proteins [43, 44]. To our knowledge, no ribosomal protein has been implicated so far in altering the levels of tRFs, but we show here that RPL24 depletion resulted in elevated levels of 5’-half tRFs. The 5’-half tRFs are produced by ANG cleavage of the anticodon loop domain of mature tRNAs [29] and are known to be elevated in response to cellular stress [8, 30]. Indeed, over expression of ANG resulted in the upregulation of 5’-half tRFs, some of which were elevated following RPL24 KD. Depleting a component of the ribosome by RPL24 KD results in translation interference [25], which would lead to cellular stress. That no direct interaction was identified between RPL24 and ANG in MS following FLAG-RPL24 IP strengthens the notion that upregulation of the 5’-half tRFs reflects a global response to the stress caused by RPL24 KD rather than a direct involvement of RPL24 in their production. Unlike the specific elevation in 5’-half tRFs observed in stressed mammalian cells, we found that mutant plants lacking the RPL24 homolog STV1 show both up- and down-regulated tRF profiles. This may reflect the difference between the acute knockdown of mammalian RPL24 and the chronic interference with STV1 function in mutant plants.

FLAG-RPL24 IP experiments provided a deeper understanding of the mechanism by which nuclear RPL24 executes its inhibiting effect on pri-miR processing. We identified a direct interaction of RPL24 with a component of the microprocessor, DDX5. In *Arabidopsis*, STV1 binding to pri-miRs promotes their processing, though STV1 was not found to interact directly with the plant microprocessor [22]. Hence, while characterizing a conserved role of RPL24 in pri-miR processing, its mechanism may be different to the one in plants. In these experiments we also identified enriched pri-miRs of those RPL24-inhibited miRs (miR-608, miR-196b), but not of RPL24-upregulated miRs (miR-185, miR-126). We conclude that RPL24 inhibition occurs upon its binding to pri-miRs in the nucleus, whereas the pri-miR sequence of miRs which are promoted by RPL24 are not bound by RPL24. This suggests that the six miRs elevated following RPL24 depletion are affected by other mechanisms such as translational repression; Three of these miRs (miR-185, miR-126 and miR-103a) are significantly downregulated by depletion of another ribosomal protein, RPS15 [45]. Interestingly, our IP results show that RPS15 interacts with RPL24 in the cytoplasm. Thus, in the cytoplasm RPL24 depletion might indirectly affect miR levels due to its interaction with other ribosomal proteins and the general effect of translational repression. Importantly, In *Arabidopsis*, STV1 indirectly influenced the transcription of many pri-miRs, and altered both the occupancy of Pol ll at those miRs’ promoters and the levels of many pri-miRs in STV1-mutant plants. It was suggested that STV1 positively or negatively affects the levels of transcription factors that regulate miR transcription by upregulating some miRs in STV1-1 mutant plants via indirect influence [22]. However, RPL24’s involvement in transcription was not tested in our study and calls for future research, as it may shed more light on additional mechanisms by which RPL24 may contribute to mammalian miR biogenesis.

miR-608, which was the initial focus of our study, targets AChE, IL-6 and the CDC42 Rho GTPase and plays a part in regulating the cholinergic blockade of inflammation [15] and stress responses [16]. Two SNPs impacting both miR-608 levels and its binding to AChE have been identified; Carriers of a SNP which resides in the miR-608 gene itself show reduced miR-608 levels, limited miR-608 regulation of AChE and other targets, and are protected from sepsis following head injury [18] and carriers of a SNP in the 3’-UTR of the AChE gene show reduced miR-608 blockade of AChE, and lowered cholinergic tone in [16]. Our current findings identify novel mechanisms regulating miR-608 levels and indicate potential functional and clinical implications to the altered levels of miR-608 in the carriers of these SNPs.

Taken together, we have identified a novel evolutionarily conserved role for RPL24 in mammalian miR biogenesis, and showed that a portion of RPL24 is located in the nucleus, where it binds pri-miRs and suppresses their expression through direct interaction with the microprocessor. Additionally, RPL24 affects the levels of 5’-half tRFs, though in an indirect manner. Our findings characterize a pan-mammalian extra-ribosomal role of RPL24 in miR biogenesis and reveal potential routes explaining the physiological impact of this phenomenon in primates.

## Acknowledgements

The research leading to these results received funding from the European Research Council under the European Union’s Seventh Framework Programme (FP/2007-2013) / ERC Grant Agreement no. 321501, to HS), The E2000 Research program led by Prof. Dr. M. Kress, Innsbruck, by the Israel Science Foundation’s Legacy Heritage Award, Grant no. 378/11 and by the ISF (1016/18; to H. Soreq). YT’s and NM’s fellowships were supported by the US friends of the Hebrew University of Jerusalem and The Sephardic Foundation on Aging (New York). KW was a Shimon Peres post-doctoral Fellow at ELSC and SD is an Azrieli PhD fellow. Parts of the figures in this article were created with BioRender.com.

## Data availability statement

The original gene expression data (FASTQ files, metadata and tables of raw counts) from the RPL24 knockdown experiments in HEK293T cells are available at the NCBI GEO database under the accession number GSE224338. All other relevant datasets have been included as supplementary files to the manuscript and the original files and code are available upon request from corresponding authors.

## Materials and Methods

### miR-608 knock-in mice

“Humanized” C57BL/6 miR-608 KI transgenic mice were produced in the EMBL Mouse Genomic Center, Monterotondo, Italy. Pre-hsa-miR-608 was inserted into the third intron of the mouse Semaphorin-4G gene (*SEMA4G*) along with 250 endogenous flanking bases on each side. The targeting vector was introduced into R1 embryonic stem (ES) cells by standard methods. Resistant ES cell colonies were screened by Southern blotting, confirmed by PCR and cells containing the modified gene were used to generate chimeric mice. One founder mouse which gave germ line transmission was bred and mice were backcrossed for at least 8 generations to minimize genetic background heterogeneity. Brain tissue from medial prefrontal cortex, hippocampus, hypothalamus, and cerebellum and peripheral tissue from liver, kidney, heart, and intestine, were obtained from wild type and transgenic mice, both female and male.

### miR-608 constructs for cell culture

miR-608 constructs were inserted into the second intron of the human hemoglobin subunit beta (*HBB*) gene and cloned into the mammalian vector pcDNA3.1+ or into the bacterial vector pUC57.

### Cell culture

Human embryonic kidney cells (HEK293T, ATTC CRL-3216) and mouse colon carcinoma cells (CT26.WT, ATTC CRL-2638) were grown under standard conditions (see Supp. Materials and Methods) and guaranteed to be free of mycoplasma. Transfection was performed using Polyethylenimine (PEI) in HEK293T cells and FuGENE™ HD Transfection Reagent (Promega, E2311) in CT26.WT cells. Cells were harvested 24 or 48 hours post-transfection.

### RNA extraction

RNA was extracted using the miRNeasy Mini Kit (Qiagen, 217004) according to the manufacturer’s protocol, followed by RNA concentration determination (NanoDrop 2000, Thermo Scientific) and standard gel electrophoresis.

### qPCR

Synthesis of cDNA from mRNA and qPCR were done using Quantabio reagents and human-specific primers. Synthesis of cDNA from microRNA and qPCR were done using either Quantabio or TaqMan™ (Thermo Fisher Scientific) reagents. The CFX384 Touch Real-Time PCR System (Bio-Rad) was used for quantification and the CFX Maestro software (Bio-Rad v4.1.2433.1219) for analysis. Data is presented as relative expression (ΔΔCt) normalized to housekeeping genes and plotted in GraphPad Prism 8.0 (GraphPad Prism Software) (See Supp. Materials and Methods).

### Core promoter

The “YAPP Eukaryotic Core Promoter Predictor” bioinformatic tool (http://www.bioinformatics.org/yapp/cgi-bin/yapp.cgi) was used to analyze the 150 bases 5’ upstream to pre-miR-608 and to confirm that the predicted TATA box could be abolished by inserting a point mutation. The TATA box was mutated by a thymine-to-adenine at position 115, using the QuikChange II site-directed mutagenesis kit (Agilent, 200521).

### Oligonucleotide pull-down assay

HEK293T cells were seeded in 100 mm plates, lysed 24 hours later, and incubated with a 5’-biotinylated ssDNA oligonucleotide (sequence identical to the 150 bases upstream to pre-miR-608). Samples were incubated in a Thermo-shaker and streptavidin beads used to retrieve the oligonucleotide together with interacting proteins (for MS analysis) and RNA (for qPCR or RNA-seq) (See Supp. Materials and Methods).

### RPL24 immunoprecipitation

HEK293T cells were seeded in 150mm plates and transfected 24 hours later with pcDNA3.1+ containing an insert of RPL24 labeled with C-terminal *Flag®-tag*) or no insert as control( and miR-608 plasmid. 48 hours later cells were lysed in buffer containing 0.1% Triton X-100 and fractionated as above. Protein concentrations were determined and both fractions were incubated with Anti-FLAG® M2 Magnetic Beads (Merck, M8823). Beads were washed once then each sample was split for MS, immunoblotting, and RNA extraction. All samples were then washed three additional times (MS samples in detergent- and glycerol-free buffer) (see supp Materials and Methods).

### Mass spectrometry

The bead-bound immunoprecipitated samples were denatured with 8M urea, treated with iodoacetamide, and trypsinized. Peptides were acidified with formic acid and desalted. MS/MS was performed on a Q Exactive™ Plus mass spectrometer coupled to a Dionex UltiMate 3000 system with peptides separated over a non-linear gradient of acetonitrile on a reverse phase C18 column. Data were acquired using Xcalibur™ software (all Thermo Fisher Scientific) and processed using the MaxQuant computational platform against a human reference proteome (UniProt UP000005640). Peptides with a length of at least seven amino acids were analyzed with FDR set at 1%. Relative protein quantification was determined using the label-free quantification algorithm. Statistical analysis was performed using the Perseus statistical package [46] with default software parameters for all statistical computations (See Supp. Materials and Methods).

### Subcellular fractionation

HEK293T cells were seeded in 6-well plates, harvested 24 hours later, and lysed in buffer containing 0.1% Triton X-100. Nuclei were pelleted by centrifugation, supernatant containing the cytoplasmic fraction was collected and the pelleted nuclei lysed. Nuclear and cytoplasmic fractions were further clarified by high-speed centrifugation (See Supp. Materials and Methods).

### Immunoblots

Protein concentrations were determined using the Bradford (Merck, B6916) or Lowry assay (DC Protein Assay, Bio-Rad, 5000113) **and** 5 μg/sample was loaded onto 4-15% gradient polyacrylamide gels (Mini-PROTEAN TGX Gels, Bio-Rad, 4561083), transferred (Bio-Rad, Trans-Blot Turbo Transfer System) to nitrocellulose membranes (Bio-Rad, 1704158) and probed with antibodies against RPL24 (Proteintech, 17082-1-AP, 1:1000), B-Actin (Santa Cruz, sc-47778, 1:1000), α-Tubulin (Merck, T5168, 1:1000), GAPDH (Cell Signaling Technology, #2118, 1:1000), Histone H3 (abcam, ab1791, 1:1000) and DDX5 (Proteintech, 10804-1-AP, 1:700).

### RPL24 knock-down

HEK293T cells were seeded and 24 hours later transfected with 50 nM of siPOOLs targeting RPL24 or non-targeting siPOOLs as control (ON-TARGETplus siRNA, Horizon, Perkin Elmer) using HiPerFect transfection reagent (301705, Qiagen). 48 hours later cells were transfected with miR-608 plasmids. After an additional 24 hours cells were harvested and RNA and protein extracted (See Supp. Materials and Methods).

### Small RNA sequencing

RNA was extracted from HEK293T cells as described above and RIN determined to be 10 for all samples (Bioanalyzer 6000, Agilent). Libraries were constructed from 800 ng total RNA (NEBNext Multiplex Small RNA library prep set for Illumina, New England Biolabs, NEB-E7560S) and the small RNA fraction was sequenced on the NextSeq 500 System (Illumina) at the Center for Genomic Technologies Facility, the Hebrew University of Jerusalem.

### Analysis of small RNA sequences

Quality control parameters in the HEK293T (above) and *Arabidopsis thaliana* (GSE91013, [22]) datasets were checked using FastQC (*http://www.bioinformatics.babraham.ac.uk/projects/fastqc/*). Reads were further trimmed and filtered using Flexbar (version 0.11.9 [47]). HEK293T sequences were aligned using miRExpress 2.1.4 [48] to miRBase version 21 for microRNAs and MINTmap 1.0 [28] for tRFs. *Arabidopsis thaliana* sequences were analyzed with the tRFanalyzer pipeline [31]. Raw files and metadata are available at the NCBI GEO database (accession number GSE224338). Differential expression analysis was performed in R version 4.0.2 using DESeq2 [27].

### Size selection for tRF qPCR

To validate sequencing results we quantified tRFs while excluding whole tRNA molecules. Total RNA samples obtained from HEK293T cells following RPL24 KD were separated by PAGE and bands containing RNAs ≤50-nt were excised from the gel. RNA was eluted, cDNA was synthesized from 500 pg/sample, and tRFs quantified with qPCR as described above (See supp Materials and Methods).

